# Regional heterothermy in *Megasoma gyas* is not related to active heat dissipation by the horns

**DOI:** 10.1101/2025.02.04.636532

**Authors:** Danilo Giacometti, Luiz Henrique Lima Silva, Guilherme Gomes, José Eduardo de Carvalho, Alexandre V. Palaoro

## Abstract

Animals rely on physiological and behavioral processes to maintain thermal balance. Some animals, however, bear structures that help dissipate excess heat when body temperatures rise. Although widespread in animals, animal weapons—exaggerated morphological structures with multiple characteristics that can make them good at dissipating heat—have rarely been studied in the context of thermoregulation. Here, we investigated whether the horns of the Rhinoceros Beetle (*Megasoma gyas*) acted as a thermal window. We heated live and dead beetles to 30ºC and allowed them to cool to 20ºC while measuring surface temperature changes in four body regions: the cephalic and thoracic horns, the scutellum, and the abdomen. If horns actively dissipated heat, they would show the lowest cooling rate among body regions. Contrary to this expectation, we found that the cephalic horn had the highest cooling rate, followed by the abdomen, thoracic horn, and scutellum, respectively. This suggests that the horns are not used for active heat dissipation in *M. gyas*. The low cooling rate of the scutellum can be explained by the presence of large flight muscles in the thorax, which play a role in heat generation, but could also aid in heat dissipation by pumping hemolymph across tagmata or through the low-insulated cuticle to prevent thoracic overheating. We also demonstrate that beetles show regional heterothermy even in the absence of exercise or stress. As such, we propose that regional heterothermy may result from both active (control of hemolymph flow) and passive (heat dissipation through poorly insulated structures) processes within individuals.

## Letter to the editor

Dear Editor,

We all know that animals possess diverse physiological and behavioral means to control body temperature (*T*_b_) (Tattersall et al., 2012). Efficient thermoregulation requires coordination between morphology and the control of internal bodily systems. A remarkable instance is found in the toucan’s beak—an exaggerated structure used to actively dissipate excess heat (Tattersall et al., 2009). However, there is another category of exaggerated morphological structures that has received little attention in the context of thermoregulation: animal weapons. Animal weapons evolved independently multiple times, possibly due to the universal need to fight for resources (Palaoro and Peixoto, 2022; Rico□Guevara and Hurme, 2019). Weapons tend to be not only large but also heavy, showing a large surface area-to-volume ratio. These two characteristics suggest that some weapons could function as thermal windows (Darnell and Munguia, 2011; Windsor et al., 2005). Evidence from the literature is scarce and mixed, but exaggerated beetle horns are good candidates for weapons that might aid in thermoregulation, since horns have a large surface area-to-volume ratio (Christiansen, 2006; Zhang et al., 2019), are not thermally isolated, and are filled with hemolymph (Shepherd et al., 2008).

While most insects exchange heat primarily through the abdomen (May, 1979), species from hot and dry habitats might use alternative routes for thermoregulation. Morphophysiological adaptations that minimize hydric, thermal, and energetic stresses experienced by individuals from hot and dry habitats have been demonstrated in both vertebrate and invertebrate taxa (Cloudsley-Thompson, 1975; Giacometti et al., 2022). In insects, the abdomen has the spiracles and a thinner cuticle than other body parts (Prange, 1996). Thus, both transcuticular and respiratory evaporative cooling can happen in the abdomen. However, if temperature and water are limiting resources, abdominal heat exchange might increase water loss, ultimately impacting the maintenance of thermal and water balance (Prange, 1996). Exchanging heat through a thicker cuticle—like the horns— might be an alternative solution to cool down while minimizing water loss.

We investigated if the horns of *Megasoma gyas* assisted in active heat dissipation. *Megasoma gyas* inhabits the Caatinga, a Brazilian semi-arid biome characterized by low rainfall and high temperatures (dos Reis Luzzi et al., 2016). Therefore, this species can be exposed to instances of thermal stress that may require active heat dissipation. We explored potential thermoregulatory roles of the cephalic and thoracic horns (Figure 1), considering that either of them might play a role in heat exchange. To assess horn contribution to heat dissipation, we compared cooling rates between live (*n* = 6) and euthanized (*n* = 3) beetles. We exposed beetles to a heat source (30ºC), allowing their *T*_b_ to passively increase for 15 min. Subsequently, we used infrared thermography to monitor surface body temperature (*T*_surface_) every 30 s for 5 min and calculate cooling rates (Dzialowski and O’Connor, 2001).

**Figure 1.**
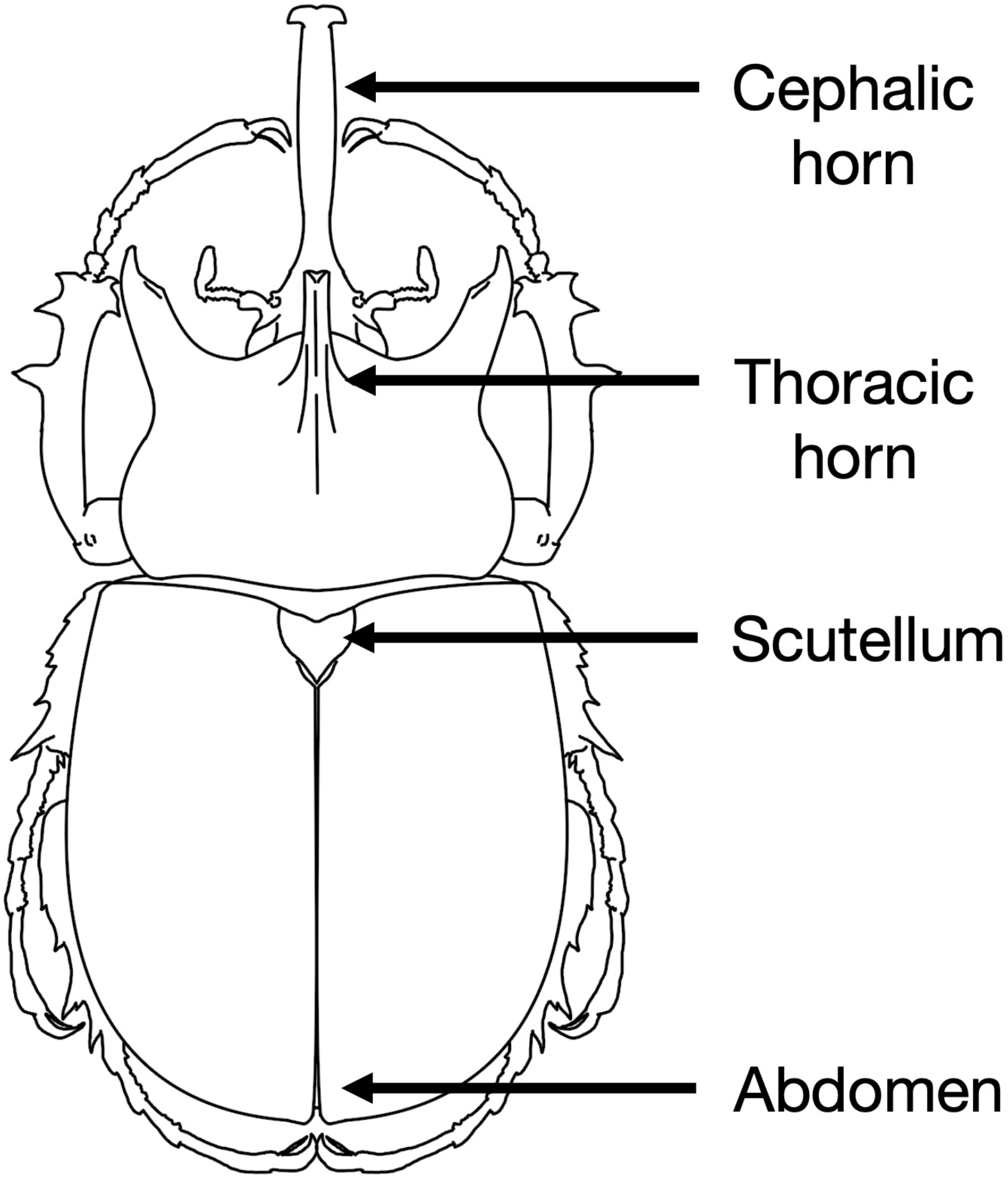
Body parts of *Megasoma gyas* from which surface temperature was measured in the current study.

Details on our procedures are available in the Supplementary Methods. If the horns function as thermal windows, we should observe higher *T*_surface_ from the horns following heat gain, as well as a lower cooling rate compared to other body parts. That is, the horns would take longer to cool down than the other body parts, since they would be actively dumping excess heat to the environment.

Contrary to this expectation, we found that the cephalic horn had the highest cooling rate, followed by the abdomen, thoracic horn, and scutellum (*F*_(3)_ = 4.573, *p* = 0.13) (Figure 2). Body mass did not mediate heat exchange (Table S1), although we must interpret these results with caution due to our limited sample size (Casey, 1988). Live beetles had lower cooling rates than control specimens for all body parts (average ± standard deviation; live specimens: abdomen = 0.22 ± 0.13, cephalic horn = 0.48 ± 0.40, scutellum = 0.07 ± 0.08, thoracic horn = 0.07 ± 0.04; control specimens: abdomen = 0.52 ± 0.44, cephalic horn = 0.74 ± 0.28, scutellum = 0.44 ± 0.48, thoracic horn = 0.68 ± 0.31). This difference was more pronounced in the thoracic horn and scutellum compared to the cephalic horn and abdomen (Figure 3). While removing the lightest individual from the analyses did not alter the qualitative interpretation of our results (Table S2), the cooling rate of the cephalic horn of this individual was two times higher than the other beetles. The lightest individual also had the smallest proportional horn size in our sample (Table S3).

**Figure 2.**
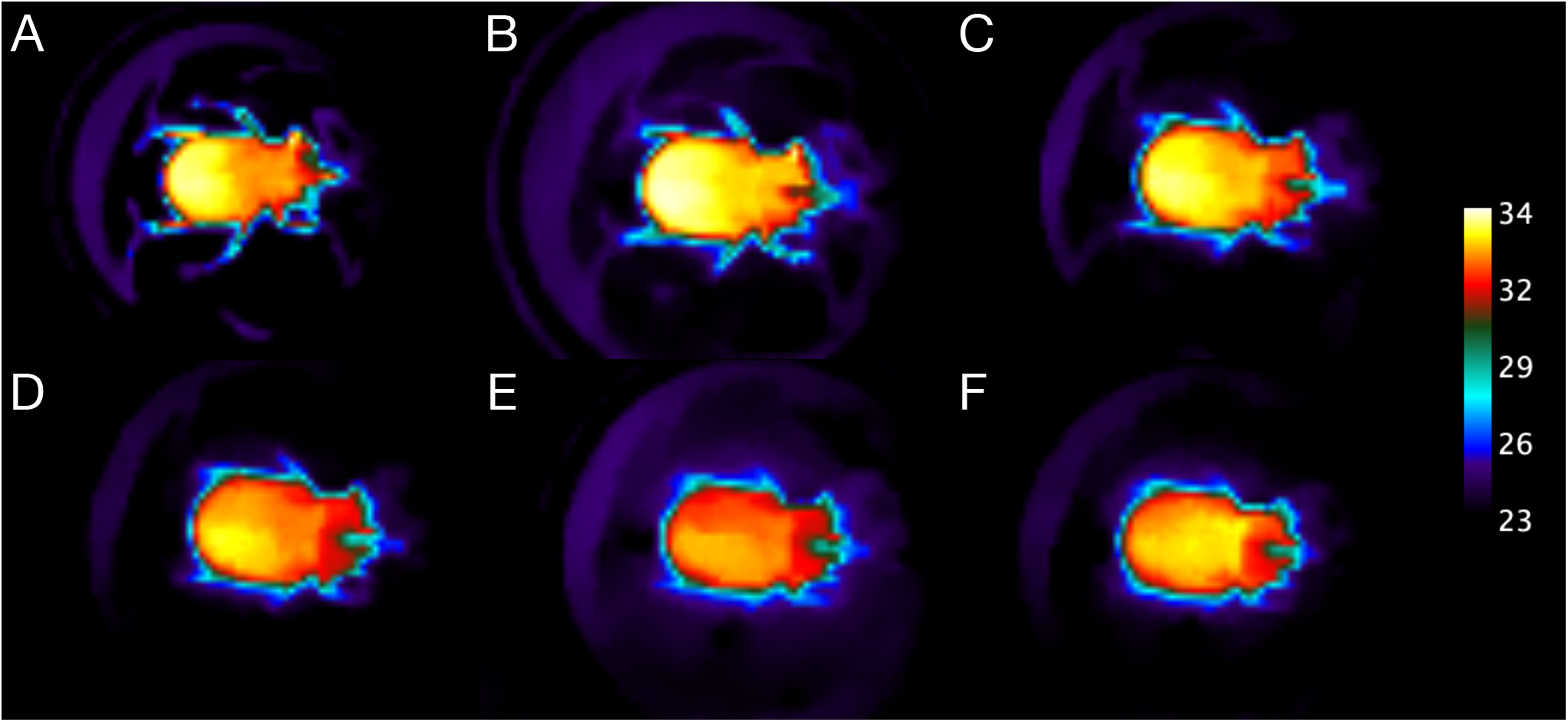
Thermal images showcasing temporal changes in heat dissipation across body parts in *Megasoma gyas*. Each letter depicts a 1-min increment from the (A) start (t = 0 min) until the (F) completion (t = 5 min) of the experiment.

**Figure 3.**
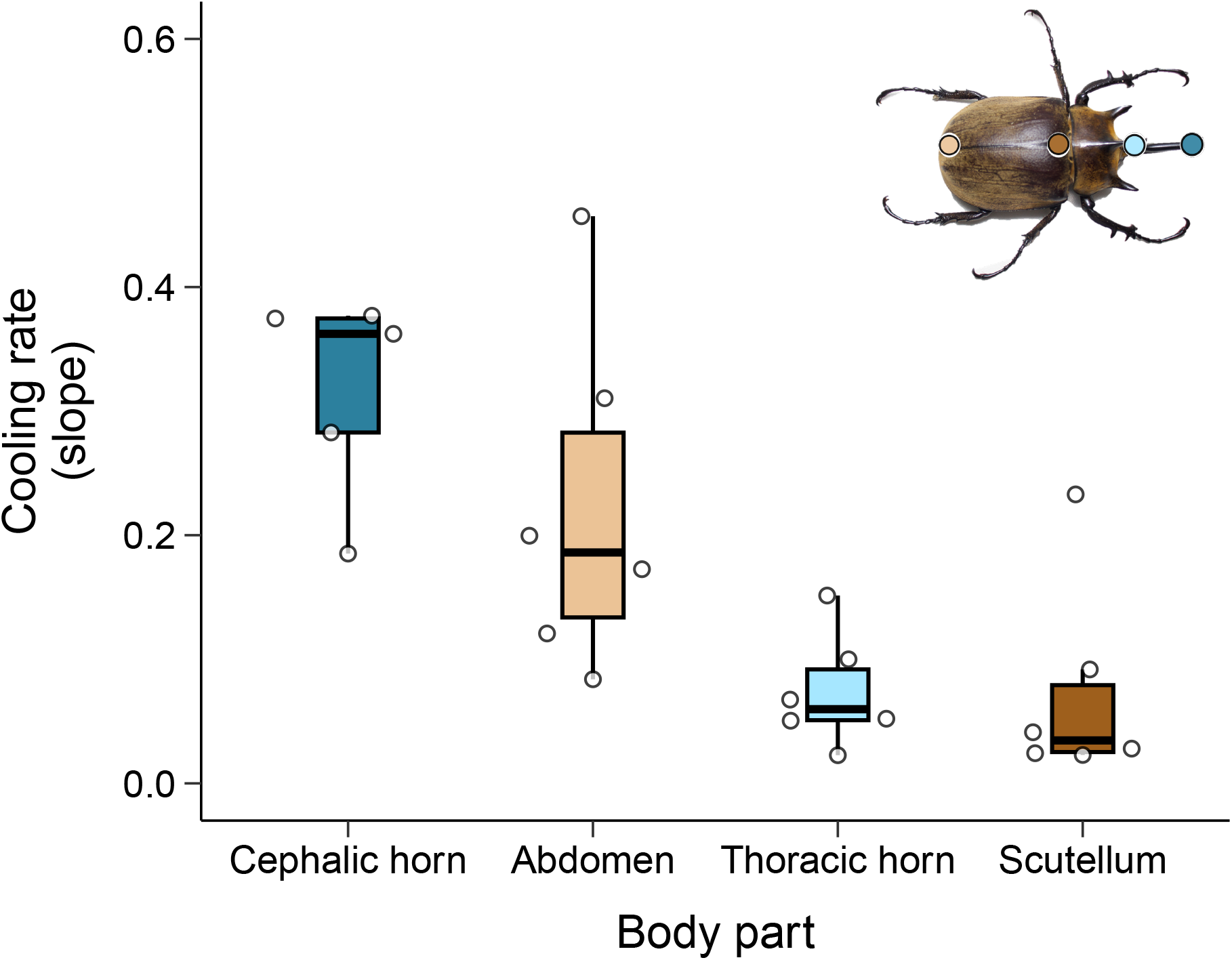
The cephalic horn and the abdomen had the highest cooling rates of the body parts measured in *Megasoma gyas*. Since body mass did not affect cooling rates, we removed it from the figure. Boxes show the 25^th^ and 75^th^ quartiles, and the horizontal line shows the median cooling rate.

Our results demonstrate that ectotherms with open circulatory systems can show regional heterothermy even when they are not performing an exercise. Regional heterothermy has been reported in both endotherms and ectotherms (Giacometti et al., 2021; Rummel et al., 2019), although its underlying mechanisms are likely manifold and context dependent. In invertebrates, regional heterothermy can occur through active (e.g., exudation of fluid drops) and /or passive processes (e.g., wind flow may cause appendages to remain cooler than the core) (Lahondère and Lazzari, 2012; Tsuji et al., 1986). A key gap that needs to be addressed is that we still do not know if regional heterothermy has a thermoregulatory role when invertebrates are not under stress or exercising (Pincebourde et al., 2013).

The cephalic horn contributed to regional heterothermy in *M. gyas*. This observation, however, does not mean that horns play a thermoregulatory role. In *Onthophagus nigriventris*, thermoregulation differed among horned males, hornless males, and hornless females. However, the possession of enlarged horns alone did not explain thermoregulatory differences between males with horns of different sizes, leading the authors to posit that sexual size dimorphism was the main effector at play (Shepherd et al., 2008). Thus, any thermoregulatory effects that the cephalic horn might have likely comes from passive mechanisms emerging from the physics of heat loss. In scarabaeid beetles, the cephalic horns are disproportionately large structures (Shepherd et al., 2008) that show a large surface area-to-volume ratio and low thermal inertia (Wang et al., 2021; Zhang et al., 2019). Hence, the horn may passively dissipate heat regardless of hemolymph flow control.

The low cooling rate of the scutellum can be explained by the large flight muscles in the thorax, which play a role in heat generation, but could also aid in heat dissipation by pumping hemolymph across tagmata or through the low-insulated cuticle to prevent thoracic overheating (Heinrich, 1993). The thorax also has a low surface-to-volume ratio and relatively high thermal inertia (May, 1979), which are associated with increased heat production and lower cooling rates. Our finding that live specimens had lower cooling rates than control specimens across all body regions could be an indication of two co-occurring mechanisms: hemolymph flow may contribute to heat dissipation, while losing water through the cuticle can decrease *T*_surface_ through convection (Gomes et al., 2018; Heinrich, 1993). Importantly, we do not know the extent to which heat can be dissipated from the core to the extremities through open circulatory systems.

In beetles, it has long been hypothesized that the thoracic pile is an inefficient insulator, and that heat flows passively across tagmata (Bartholomew and Heinrich, 1978; Morgan, 1987). However, evidence suggests that some species may modulate heat transfer across tagmata when faced with high ambient temperatures (Verdú et al., 2004). Our finding of regional heterothermy in the absence of exercise supports this notion, and we propose that regional heterothermy may result from both active (hemolymph flow) and passive (heat dissipation through poorly insulated structures) processes. We acknowledge that our sample size is limited, and the maximum temperature we used in our study might not have been sufficient to actively move excess heat (Vorhees and Bradley, 2012). However, as far as we know, this is the first report of regional heterothermy in these giant insects that naturally occur in a neotropical semi-arid biome.

## Acknowledgements

We thank Dr. Glenn Tattersall for helpful comments about the heat exchange dynamics of regional heterothermy and guidance on thermal image analysis. We also thank Dr. Sônia Casarin, the Coleoptera curator at MZUSP, and Dr. Juares Fuhrmann, for lending us the beetles (both live and dead) and all the help provided. DG was funded by a Doris White Memorial Bursary provided by Brock University. JEC was supported by the Fundação de Amparo à Pesquisa do Estado de São Paulo (FAPESP; grant: 20 / 12962-5).

