## Supplementary methods for "Regional heterothermy in *Megasoma gyas* is not related to active heat dissipation by the horns"

*Animal collection and husbandry*

Six live adult male *M*. *gyas* were rescued from animal trafficking at Guarulhos International Airport (State of São Paulo, Brazil) and transported to the Museum of Zoology at the University of São Paulo (MZUSP) (Leite, Isabela. São Paulo Customs seized 99 live beetles in passenger's suitcase bound for Thailand. 2020. Available in Portuguese at: <https://g1.globo.com/sp/sao-paulo/noticia/2020/02/29/receita-de-sp-apreende-99-besouros-vivos-em-mala-de-passageiro-que-iria-a-tailandia.ghtml>). We were granted access to the *M. gyas* individuals for use in our experiments before they were euthanized and added to the MZUSP collection. Prior to the experiments, we individually housed each specimen in a plastic container (3L) with sawdust and a piece of banana, allowing them to habituate for one day in the laboratory at 20°C. To establish the baseline of how much heat exchange occurred through the cuticle we used three pinned (dead; hereafter “control”) beetles from the MZUSP collection. All individuals were returned to the MZUSP afterwards.

*Measuring* T*_surface_*

Our goal with the experiment was to raise the *T*_b_ of the individuals and analyze how they exchanged heat with the environment until returning to the initial temperature of the experiment. To measure *T*_b_, we used a thermal imaging camera (FLIR C-3, Wilsonville, OR, USA; image resolution: 128 x 96 pixels, temperature measurement accuracy: ± 2°C). Thermography is suitable for measuring *T*_surface_, because it allows for non-invasive and instantaneous temperature recording (Tattersall, 2016). Furthermore, thermography also enables us to capture images of the individuals over time, facilitating the analysis of temperature exchange patterns between the focal object and the surrounding environment continuously. Finally, thermography allows us to measure the *T*_surface_ of various body parts simultaneously, enabling us to compare how different body parts exchange heat. As such, thermography is ideal for answering our question about the horn's capacity to serve as a thermal window.

*Passive heating*

To analyze heat exchange, we conducted a passive heating experiment with live and control individuals. We recorded the body mass of the six live individuals and measured their resting *T*_surface_ using thermography before starting the experiment. Then, we placed each individual in a glass container that was partially submerged in a constant-temperature water bath set to 30°C for 15 minutes. Therefore, the heat source for this treatment was external and aimed at simulating the exposure to a relatively high ambient temperature that may occur in nature while heating the animal uniformly throughout its volume. After the water bath, we recorded *T*_surface_ every 30 seconds for five minutes with the thermal camera. Finally, we recorded the body mass of the individuals again at the end of the experiment and returned them to their respective containers. We used the same procedure for the control individuals, with the only difference being that, in addition to the thermal camera, we also used a thermocouple (TC-200 Type T Thermocouple Meter – Sable Systems International) to calibrate the absolute temperatures of the thermographs.

*Thermal image processing*

We processed all thermal images using ThermimageJ (Tattersall, 2019) plugins for Fiji (ImageJ) (Schindelin et al., 2012) that were verified against algorithms in FLIR Tools software (FLIR Systems, Wilsonville, OR, USA). We assumed an object emissivity of 0.95 (Tattersall, 2016) and adjusted parameters (e.g.*,* object distance, atmospheric temperature, reflected temperature) according to laboratory conditions recorded on the day of the experiment. For each thermal image, we obtained measures of *T*_surface_ by drawing a region of interest (ROI) over four body regions: (I) cephalic horn, (II) central pronotal horn (thoracic horn), (III) scutellum, and (IV) abdomen (Figure 1). We selected these regions of the beetle's body because: (1) the horns are the weapons we are interested in testing our hypothesis on; (2) the scutellum is a region where there is a high concentration of flight muscles, and thus is a region that can generate heat (Morgan, 1987); and (3) the abdomen is another region that could aid in heat exchange (Heinrich, 1974). To correct for possible deviations in the absolute temperatures measured with thermography, we measured the temperature of the spot the animals were placed as a control measure of room temperature, since room temperature should directly and dynamically influence *T*_b_ during the procedures. Additionally, in the procedure with the control individuals, we used a thermocouple to measure room temperature. The substrate temperature measurement collected from the thermocouple is more accurate than that obtained from thermography. Thus, by comparing the two measurements, possible differences between thermography and thermocouple measurements can be corrected.

*Measuring body and horn size*

After the experiments, we imaged the individuals to measure both their body and horn size. We positioned all individuals on a white flat surface with a ruler next to them. The camera was then positioned on top the surface and all beetles were imaged from their lateral side. That position guaranteed that the camera was aligned with the axis of the individual (i.e., not tilted), while also allowing us to measure horn length while accounting for the natural upward curvature of the horn.

We used ImageJ (Schneider et al., 2012) to measure the individuals. Body size was measured as the length of the elytrum, measured from its insertion on the thorax to the distalmost point on the abdomen. Horn size was measured using a segmented line, where we positioned the line along the dorsal side of the cephalic horn to encompass its natural curvature. We then divided horn length by body length to obtain a measure of proportional horn size for each beetle.

*Statistical analyses*

We performed all analyses with RStudio (version 2023.06.2) in R (version 4.2.2) (R Core Team, 2024), with a significance level of 0.05. To test our prediction that the horns would have a lower cooling rate than other body parts, we first calculated the cooling rate of each body region for each individual. To do this, we calculated a linear relationship between the surface *T*_b_ of each body part and time. The slope of this relationship indicates how much the temperature decreases per unit of time—in other words, the cooling rate of the respective region (Dzialowski & O’Connor, 2001). The curve of this cooling rate is decreasing (as temperature decreases over time), but to facilitate the analysis and interpretation of the results, the values were multiplied by -1. This way, the curve becomes positive, and the higher the cooling rate (i.e., the steeper the regression slope), the more heat was dissipated per unit of time. Additionally, heat exchange with the environment is also influenced by the size of the individuals, which required us to control for body mass when testing our hypothesis.

To test our hypothesis that the horns acted as thermal windows, we used the cooling rate as the dependent variable, the body mass of the individual, and the body part as independent variables in a linear model fit with the ‘lm’ function from the “stats” package (R Core Team, 2024). We did not include an interaction between the independent variables because we used body mass only to control the effect of the size of the individual. Since body mass did not explain cooling rates, we report the results of a model fit with the ‘aov’ function from the “stats” package (R Core Team, 2024) which had cooling rate as the dependent variable and body parts as independent variables. It is worth acknowledging that we capitalized on the opportunity to work with a species that is extremely hard to find in nature (dos Reis Luzzi et al., 2016), and we were only able to do so because our study individuals were rescued from animal trafficking. As such, we recognize that due to the relatively small sample size available for our study, we did not have the statistical power to control for pseudoreplication in our model. To validate our results, we also performed sensitivity analysis to understand whether removing the individual with the lowest body mass in our data set would affect the interpretation of our results. We assessed model assumptions with the ‘check_model’ function from the “performance” package (Lüdecke et al., 2021) and residual autocorrelation with the ‘checkresiduals’ function from the “forecast” package (Hyndman et al., 2020). To create figures, we used the “ggplot2” package (Wickham, 2016). The data and code necessary to reproduce our analyses are available in the Supplementary Material.

**Supplementary material**

**SM1.** The data and code necessary to reproduce our analyses can be accessed from: <https://github.com/alexandrepalaoro/beetle-heat>

**Supplementary tables**

**Table S1.** Parameter estimates (β) and standard error (SE) for the linear model testing how cooling rates differed among body parts while accounting for body mass in *Megasoma gyas*. P-values in bold denote significant terms in the model.

| Parameter | β | SE | t-value | P |
| --- | --- | --- | --- | --- |
| Body part [Abdomen] | 0.68 | 0.26 | 2.60 | **0.01** |
| Body part [Cephalic horn] | 0.25 | 0.12 | 2.13 | **0.04** |
| Body part [Scutellum] | -0.15 | 0.12 | -1.25 | 0.22 |
| Body part [Thoracic horn] | -0.15 | 0.12 | -1.26 | 0.22 |
| Body mass | -0.02 | 0.01 | -1.84 | 0.08 |

**Table S2.** Parameter estimates (β) and standard error (SE) for the linear model testing how cooling rates differed among body parts while accounting for body mass in *Megasoma gyas*. P-values in bold denote significant terms in the model. We removed the smallest individual from this model to understand if a potential outlier affected our results.

| Parameter | β | SE | t-value | P |
| --- | --- | --- | --- | --- |
| Body part [Abdomen] | 0.12 | 0.29 | 0.41 | 0.68 |
| Body part [Cephalic horn] | 0.07 | 0.06 | 1.15 | 0.26 |
| Body part [Thoracic horn] | -0.18 | 0.06 | -2.99 | **0.01** |
| Body part [Scutellum] | -0.17 | 0.06 | -2.81 | **0.01** |
| Body mass | 0.01 | 0.01 | 0.41 | 0.68 |

**Table S3.** Body size parameters for *Megasoma gyas*. Body mass values are reported in grams (g), whereas those for the elytrum and the cephalic horn are given in millimeters (mm). The column ‘Proportion’ represents the proportional length of the cephalic horn relative to the length of the elytrum.

| Individual | Body mass | Elytrum | Cephalic horn | Proportion |
| --- | --- | --- | --- | --- |
| 1 | 23.79 | 61.27 | 58.45 | 0.95 |
| 2 | 22.90 | 73.14 | 49.75 | 0.68 |
| 3 | 19.13 | 64.02 | 46.54 | 0.73 |
| 4 | 23.32 | 56.70 | 37.13 | 0.65 |
| 5 | 13.74 | 43.84 | 18.51 | 0.42 |
| 6 | 22.86 | 49.23 | 39.13 | 0.79 |
